# Evolutionary assembly patterns of prokaryotic genomes

**DOI:** 10.1101/027649

**Authors:** Maximilian O. Press, Christine Queitsch, Elhanan Borenstein

## Abstract

Evolutionary innovation must occur in the context of some genomic background, which limits available evolutionary paths. For example, protein evolution by sequence substitution is constrained by epistasis between residues. In prokaryotes, evolutionary innovation frequently happens by macrogenomic events such as horizontal gene transfer (HGT). Previous work has suggested that HGT can be influenced by ancestral genomic content, yet the extent of such gene-level constraints has not yet been systematically characterized. Here, we evaluated the evolutionary impact of such constraints in prokaryotes, using probabilistic ancestral reconstructions from 634 extant prokaryotic genomes and a novel framework for detecting evolutionary constraints on HGT events. We identified 8,228 directional dependencies between genes, and demonstrated that many such dependencies reflect known functional relationships, including, for example, evolutionary dependencies of the photosynthetic enzyme RuBisCO. Modeling all dependencies as a network, we adapted an approach from graph theory to establish chronological precedence in the acquisition of different genomic functions. Specifically, we demonstrated that specific functions tend to be gained sequentially, suggesting that evolution in prokaryotes is governed by functional assembly patterns. Finally, we showed that these dependencies are universal rather than clade-specific and are often sufficient for predicting whether or not a given ancestral genome will acquire specific genes. Combined, our results indicate that evolutionary innovation via HGT is profoundly constrained by epistasis and historical contingency, similar to the evolution of proteins and phenotypic characters, and suggest that the emergence of specific metabolic and pathological phenotypes in prokaryotes can be predictable from current genomes.

## INTRODUCTION

A fundamental question in evolutionary biology is how present circumstances affect future adaptation and phenotypic change (Gould and Lewontin 1979). Studies of specific proteins, for example, indicate that epistasis between sequence residues limits accessible evolutionary trajectories and thereby renders certain adaptive paths more likely than others (Weinreich et al. 2006; Gong et al. 2013; de Visser and Krug 2014; Harms and Thornton 2014). Similarly, both phenotypic characters (Ord and Summers 2015) and specific genetic adaptations (Christin et al. 2015; Conte et al. 2012) show strong evidence of parallel evolution rather than convergent evolution. That is, a given adaptation is more likely to repeat in closely related organisms than in distantly related ones. This inverse relationship between the repeatability of evolution and taxonomic distance implies a strong effect of lineage-specific contingency on evolution, also potentially mediated by epistasis (Orr 2005).

Such observations suggest that genetic adaptation is often highly constrained and that the present state of an evolving system can impact future evolution. Yet, the studies above are limited to small datasets and specific genetic pathways, and a more principled understanding of the rules by which future evolutionary trajectories are governed by the present state of the system is still lacking. For example, it is not known whether such adaptive constraints are a feature of genome-scale evolution or whether they are limited to finer scales. Moreover, the mechanisms that underlie observed constraints are often completely unknown. Addressing these questions is clearly valuable for obtaining a more complete theory of evolutionary biology, but more pressingly, is essential for tackling a variety of practical concerns including our ability to combat evolving infectious diseases or engineer complex biological systems.

Here, we address this challenge by analyzing horizontal gene transfer (HGT) in prokaryotes. HGT is an ideal system to systematically study genome-wide evolutionary constraints because it involves gene-level innovation, occurs at very high rates relative to sequence substitution (Nowell et al. 2014; Puigbò et al. 2014), and is a principal source of evolutionary novelty in prokaryotes (Gogarten et al. 2002; Jain et al. 2003; Lerat et al. 2005; Puigbò et al. 2014). Clearly, many or most acquired genes are rapidly lost due to fitness costs (van Passel et al. 2008; Baltrus 2013; Soucy et al. 2015), indicating that genes retained in the long term are likely to provide a selective advantage. Moreover, not all genes are equally transferrable (Jain et al. 1999; Sorek et al. 2007; Cohen et al. 2011), and not all species are equally receptive to the same genes (Smillie et al. 2011; Soucy et al. 2015). However, differences in HGT among species have been attributed not only to ecology (Smillie et al. 2011) or to phylogenetic constraints (Nowell et al. 2014; Popa et al. 2011), but also to interactions with the host genome (Jain et al. 1999; Cohen et al. 2011; Popa et al. 2011). Indeed, studies involving single genes or single species support the influence of genome content on the acquisition and retention of transferred genes (Pal et al. 2005; Iwasaki and Takagi 2009; Chen et al. 2011; Press et al. 2013; Sorek et al. 2007; Johnson and Grossman 2014). For example, it has been demonstrated that the presence of specific genes facilitates integration of others into genetic networks (Chen et al. 2011), and that genes are more commonly gained in genomes already containing metabolic genes in the same pathway (Pal et al. 2005; Iwasaki and Takagi 2009). However, to date, a systematic, large-scale analysis of such dependencies has not been presented. In this paper, we therefore set out to characterize a comprehensive collection of genome-wide HGT-based dependencies among prokaryotic genes, analyze the obtained set of epistatic interactions, and identify patterns in the evolution of prokaryotic genomes.

## RESULTS

### PGCE Inference

We first set out to detect pairs of genes for which the presence of one gene in the genome promotes the gain of the other gene (though not necessarily *vice versa*) (Figure 1). Such “pairs of genes with conjugated evolution” (PGCEs) represent putative epistatic interactions at the gene level and may guide genome evolution. To this end, we obtained a collection of 634 prokaryotic genomes annotated by KEGG (Kanehisa et al. 2012) and linked through a curated phylogeny (Dehal et al. 2010). For each of the 5801 genes that varied in presence across these genomes, we reconstructed the probability of this gene’s presence or absence on each branch of the phylogenetic tree using a previously introduced method (Cohen and Pupko 2010), as well as the probability that it was gained or lost along these branches using a simple heuristic (Methods). We confirmed that genes’ presence/absence was robust to the reconstruction method employed (99.5% agreement between reconstruction methods used; Methods). As expected (Mira et al. 2001), gene loss was more common than gene gain for most genes (Supplemental Figure S1, Supplemental Text). We additionally confirmed that inferred gains of several genes of interest were consistent with gains inferred by an alternative HGT inference method (Methods; Supplemental Text, Supplemental Table S1). From the reconstructions, we estimated the frequency with which each gene was gained in the presence of each other gene, and followed previous studies (Maddison 1990; Cohen et al. 2012) in using parametric bootstrapping (Supplemental Figure S2) to detect PGCEs - gene pairs for which one gene is gained significantly more often in the presence of the other (Supplemental Figure S3, Supplemental Text). In total, we identified 8,415 PGCEs. We finally applied a transitive reduction procedure to discard potentially spurious PGCEs, resulting in a final network containing 8,228 PGCEs connecting a total of 2,260 genes (Supplemental Figures S4, S5, Supplemental Text). A detailed description of the procedures used can be found in Methods, and the final list of PGCEs is supplied as Supplemental File S1.

**Figure 1.**
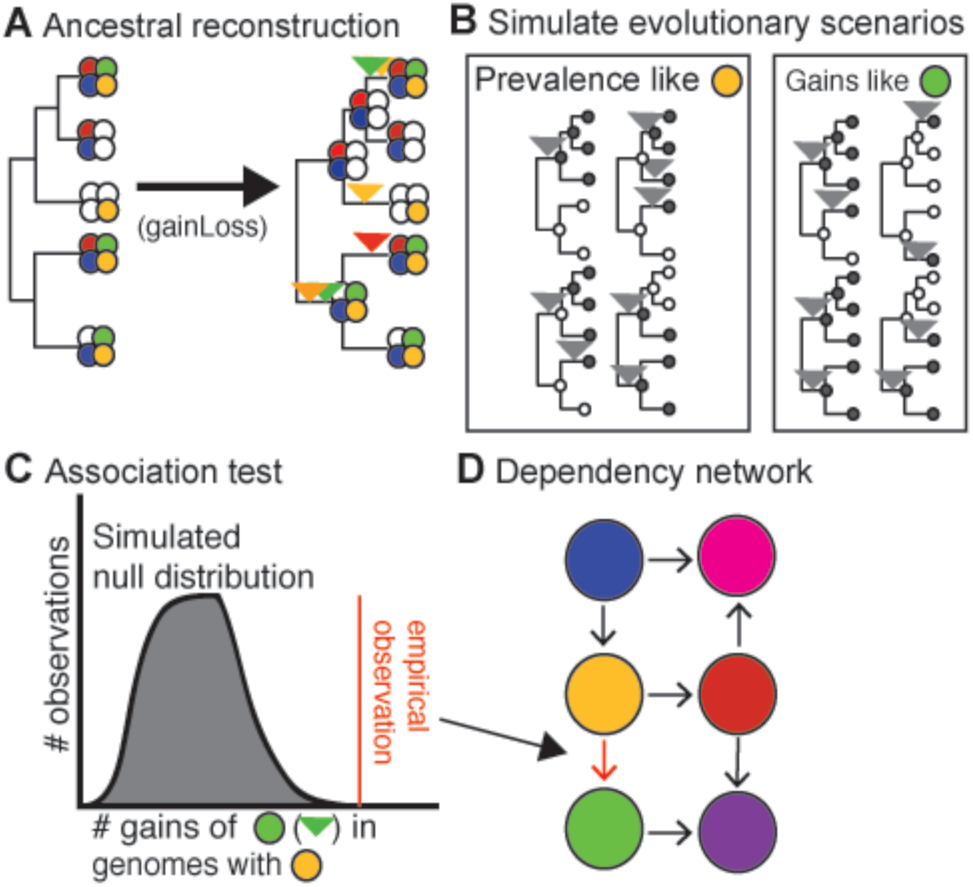
Workflow for deriving the PGCE network. (A): a model phylogeny and a set of gene presence/absence patterns at the tips are used to generate an ancestral reconstruction, from which gains are inferred. Filled circles represent the presence of a gene (distinguished by color), empty circles represent absence of that gene. Inverted triangles represent points on the phylogeny where the gene of the indicated color is inferred to be gained. (B): Based on inferred gain and loss rates, many evolutionary scenarios are independently simulated and used as a null expectation for evolutionary independence. Filled circles indicate presence of the simulated gene and empty circles indicate absence, inverted triangles represent gains of the simulated gene on the phylogeny. (C): A null distribution derived from simulated gene evolution is used to identify dependencies between real genes. (D): These dependencies are modeled as a network. Filled circles indicate genes (nodes), arrows indicate dependencies (edges).

### PGCEs represent biologically relevant dependencies

Comparing this final set of PGCEs to known biological interactions, we confirmed that the obtained PGCEs represent plausible biological dependencies. For example, genes sharing the same KEGG Pathway annotations were more likely to form a PGCE (Figure 2A), as were genes linked in an independently-derived network of bacterial metabolism (Levy and Borenstein 2013) (Figure 2B). Moreover, PGCEs often linked genes in functionally related pathways (Supplemental Figure S6, Supplemental Text). We similarly identified specific examples in which PGCEs connected pairs of genes with well-described functional relationships. One such example is the PGCE connecting *rbsL* and *rbsS* (sometimes written *rbcL/rbcS*), two genes that encode the large and small subunits of the well-described photosynthetic enzyme ribulose-1-5-bisphosphate carboxylase-oxygenase (RuBisCO), respectively. The rbsL subunit alone has carboxylation activity in some bacteria, but the addition of rbsS increases enzymatic efficiency, consistent with its PGCE dependency on *rbsL* (Figure 3A) (Andersson and Backlund 2008). Moreover, these genes are known to undergo substantial horizontal transfer (Delwiche and Palmer 1996).

**Figure 2.**
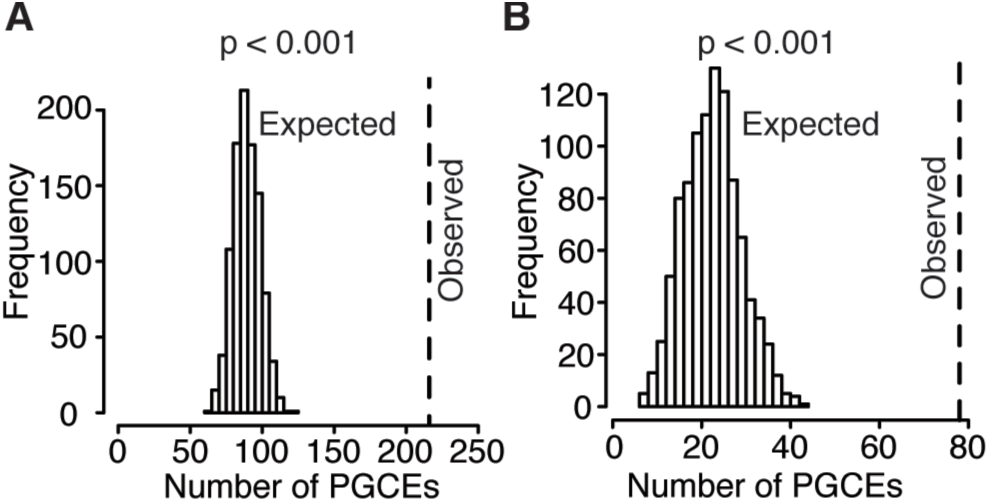
PGCEs are enriched for biologically meaningful interactions. (A): The observed number of PGCE edges connecting genes in the same pathway (dotted line), compared to the expected distribution obtained from 1000 rewired networks with identical degree distributions. (B): The observed number of PGCE edges that also appear in a bacteria-wide metabolic network, compared to the expected distribution.

**Figure 3.**
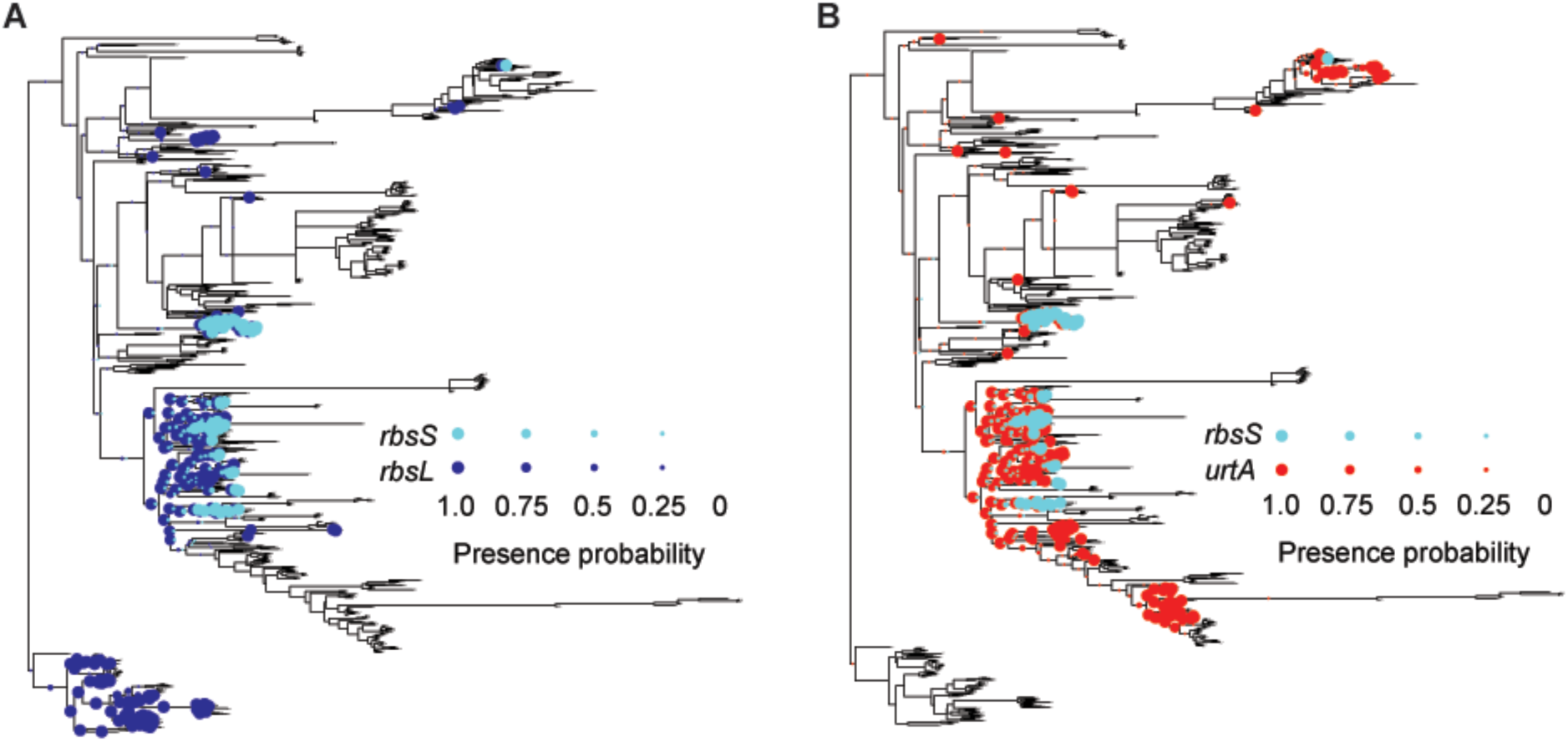
The phylogenetic history of *rbsL, urtA* and *rbsS*. The presence of each gene in each branch in the phylogenetic tree is illustrated with a colored circle, with the circle’s diameter scaled to denote the probability of presence. (A): *rbsL* and *rbsS* evolutionary histories; (B): *urtA* and *rbsS* evolutionary histories. The long branch leading to Archaea (bottom-most clade) was reduced in size for graphical purposes.

Multiple additional genes were found to promote *rbsS* gain (88 PGCEs in total, Supplemental Table S2), many of which, as expected, are associated with carbon metabolism. Other genes in this set, however, unexpectedly implicated nitrogen acquisition, as well as other pathways (Supplemental Table S3), in promoting *rbsS* gain. For example, all components of the *urt* urea transport complex had a PGCE link with *rbsS*, as shown by the reconstructed phylogenetic history of *urtA* and *rbsS* (Figure 3B). This strict dependency could reflect nitrogen’s role as a rate-limiting resource for primary production in phytoplankton and other photosynthetic organisms (Eppley and Peterson 1979; Sohm et al. 2011). In comparing the reconstructions from which *urtA-rbsS* and *rbsL-rbsS* dependencies were inferred, we further observed that *rbsS* is gained only in lineages where both *urtA* and *rbsL* were previously present. This indicates that while both *rbsL* and *urtA* may be necessary for the acquisition of *rbsS*, neither *rbsL* nor *urtA* are independently sufficient for the acquisition of *rbsS*. Other PGCEs may interact in similarly complex fashions in controlling the acquisition of genes, and thus such relationships may be gene-specific and involve a variety of biological mechanisms that may be difficult to generalize. For further analyses, we therefore focused on analyzing large-scale patterns of PGCE connectivity and on exploring how the dependencies between various genes structure the relationships between functional pathways.

### PGCE network analyses reveal evolutionary assembly patterns

The *rbsS*-associated PGCEs described above show how PGCEs captured an assembly pattern involving multiple pathways. Therefore, we next set out to infer global evolutionary assembly patterns based on the complete set of PGCEs identified. Specifically, we used a network-based topological sorting approach (Supplemental Text) to rank all genes in the PGCE network. According to this procedure, genes without dependencies occupy the first rank, genes in the second rank have PGCE dependencies only on first rank genes, genes in the third rank have dependencies only on first and second rank genes, and so on until all genes are associated with some rank. In other words, the obtained ranking represents general patterns in the order by which genes are gained throughout evolution, with the gain of higher-ranked genes succeeding the presence of the lower-ranked genes on which they depend. Using this approach, we found that genes could be fully classified into five ranks (Fig 4A). The first rank was by far the largest at 1,593 genes (most genes do not have detectable dependencies), the second rank had 498 genes, and successive ranks showed declining membership until the last (fifth) rank, with only 5 genes (Supplemental Table S4).

**Figure 4.**
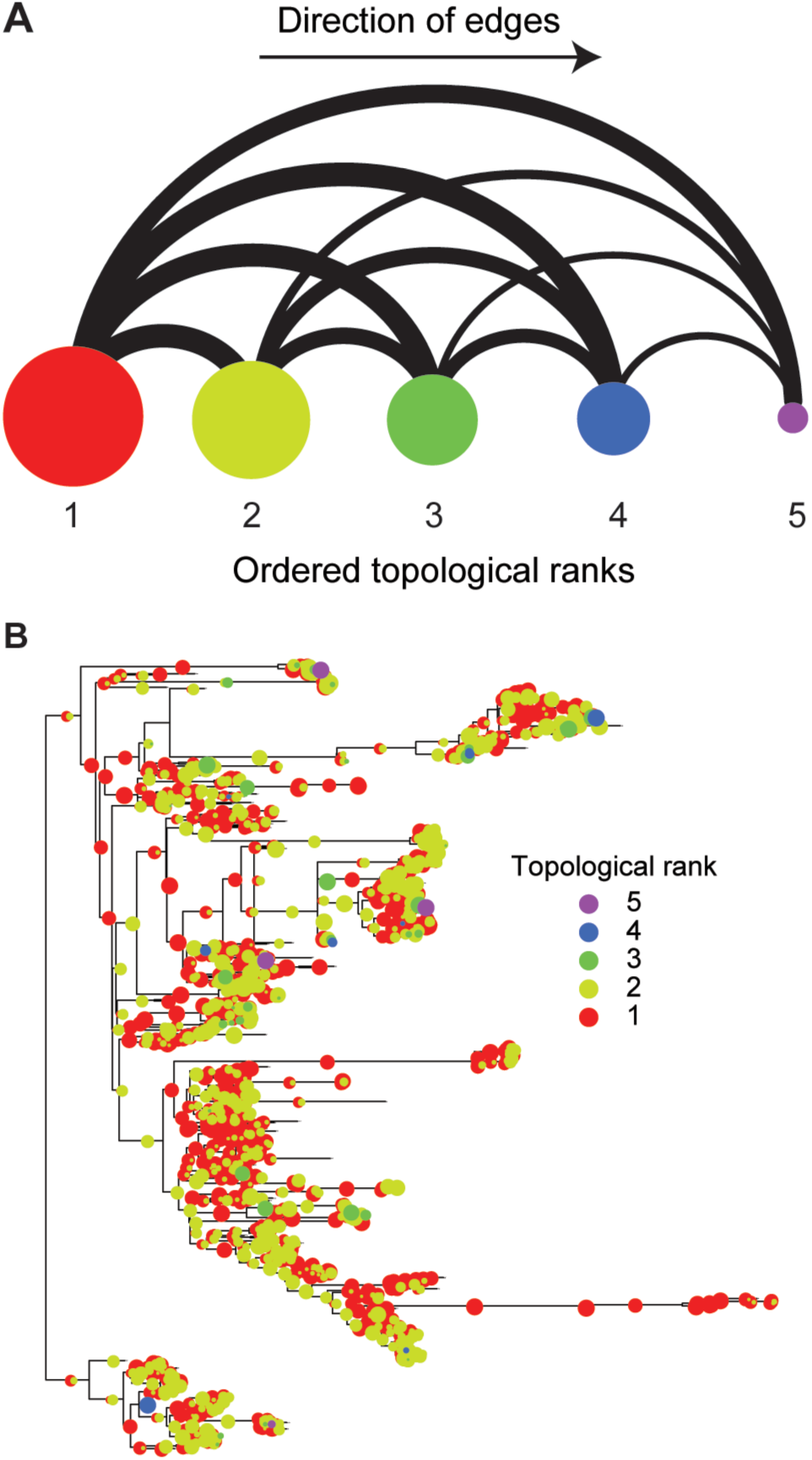
Topological sorting of the PGCE dependency network reveals assembly patterns that govern the evolutionary process. (A): Binned dependencies among the six ranks of genes in the topological sort (left to right). Node size represents the number of genes in each rank (using natural logarithm-scale). Edge width represents the number of PGCEs between genes in different rank (natural logarithm-scale), all edges are directed to the right. (B) The gain of genes from each rank in each branch of the phylogenetic tree is illustrated (circles). The different colors represent different ranks. Circle sizes correspond to the proportion of gains on a branch attributed to genes of that rank (e.g. a large red circle indicates that most gains on a branch correspond to rank 1). The branch to Archaea (lower clade) has been reduced in size for graphical purposes. See also Supplemental Figure S7.

To identify evolutionary assembly patterns from these ranks, we examined the set of genes in each rank and identified overrepresented functional categories (Table 1). These enriched functional categories indicate that certain functional groups of genes consistently occupy specific positions in these evolutionary assembly patterns, whether in controlling other genes’ gain or in being controlled by other genes. For example, we found that the first rank was enriched for flagellar and pillar genes involved in motility, in addition to Type II secretion genes (many of which are homologous to or overlap with genes encoding pillar proteins) and certain two-component genes. The second rank was enriched for various metabolic processes, whereas later ranks were enriched for Type III and Type IV secretion systems and conjugation genes. This finding suggests that habitat commitments are made early in evolution, mediated by motility genes that could underlie the choice and establishment of physical environments. This environmental choice is followed by a metabolic commitment to exploiting the new habitat. Last, genes for interaction with the biotic complement of these habitats are gained, and replaced frequently in response to evolving challenges. Considering two distinct but highly homologous pilus assembly pathways, one (fimbrial) was enriched in a low rank and one (conjugal) was enriched in a high rank, suggesting that the specific function of the gene rather than other sequence-level gene properties drove the ranking (Supplemental Figure S7A). We additionally confirmed that the observed rank distribution for these functions is not explained by variation in the frequency of gene gain (Supplemental Figure S7B). Furthermore, as expected, we observed that the gains of genes appearing late in the sort were overrepresented in later branches of the tree compared to the gains of lower-ranked genes (Figure 4B, Supplemental Figure S8), suggesting that the chronology of gene acquisition reflects the overall assembly patterns in gain order.

**Table 1.** Functional groups are enriched in different ranks of the topological sort.

| Annotation label | P-value<sup>1</sup> | Enrichment Ratio<sup>2</sup> |
| --- | --- | --- |
| **Rank 1 Enrichments** |  |  |
| Cell motility | 1.94E-07 | 1.40 |
| Bacterial motility proteins | 1.85E-11 | 1.41 |
| Type II secretion system | 2.61E-05 | 1.33 |
| Two-component system | 3.65E-04 | 1.25 |
| Flagellar system | 1.01E-09 | 1.43 |
| Pilus system | 2.11E-04 | 1.38 |
| Metabolism<sup>3</sup> | 3.37E-05 | 0.91 |
| Xenobiotics biodegradation and metabolism<sup>3</sup> | 1.07E-06 | 0.69 |
| Carbohydrate metabolism<sup>3</sup> | 0.00012 | 0.84 |
| Type IV secretion system<sup>3</sup> | 1.26E-09 | 0.20 |
| **Rank 2 Enrichments** |  |  |
| Metabolism | 1.47E-04 | 1.23 |
| Carbohydrate metabolism | 3.08E-06 | 1.58 |
| **Rank 4 Enrichments** |  |  |
| Pathogenicity | 1.88E-06 | 21.6 |
| Conjugal transfer pilus assembly protein | 1.08E-04 | 15.0 |
| Type III protein secretion pathway protein | 1.88E-06 | 21.6 |
| ABC-2 type and other transporters | 2.31E-04 | 12.5 |
| Type IV secretion system | 1.30E-03 | 8.04 |

1: from a hypergeometric test. All annotations displayed are significant at a 1% false discovery rate.
2: The ratio of the observed proportion of genes with this label in the indicated rank to the expected proportion based on all genes in the network.
3: These annotations are depleted (i.e. enrichment ratio significantly less than one) in the first rank.

### Evolution by HGT is predictable

The chronological ordering of ranks was relatively consistent across the tree (Figure 4B), indicating that PGCE dependencies are universal across prokaryotes. Notably, this universality also implies that gene acquisition is predictable from genome content. Put differently, if PGCEs are universal, then PGCEs inferred in one clade of the tree are informative in making predictions about gene acquisition in a different clade. Indeed, studies of epistasis-mediated protein evolution indicate that the constriction of possible mutational paths should lead to predictability in evolution, if epistasis is sufficiently strong (Weinreich et al. 2006). To explore this hypothesis explicitly, we partitioned the tree into training and test sets (Figure 5A). As test sets, we selected the Firmicutes phylum, and the Alphaproteobacteria/Betaproteobacteria subphyla. Choosing whole clades as test sets (rather than randomly sampling species from throughout the tree) guarantees that true predictions are based on universal PGCEs, rather than clade-specific PGCEs. For each test set, we used a model phylogeny that excluded the test subtree as a training set, and inferred PGCEs based on this pruned tree (Supplemental Table S5, Supplemental Figure S9A). We then used these inferred PGCEs to score the relative likelihood of the gain of dependent genes on each branch in the test set, based on the genome content of the branch’s ancestor (Figure 5A, Supplemental Table S5, Supplemental Text). We used a naïve and simplistic score: the proportion of genes upon which the gained gene depends that are present in the reconstructed ancestor of each branch. In both test sets, we found that prediction quality was surprisingly high (Figure 5B, Supplemental Figure S9B-C), suggesting that PGCEs are taxonomically universal and statistically robust in describing relationships between genes. This predictability is consistent with the hypothesis that gene-gene dependencies constrain the evolution of genomes by HGT. More broadly, this analysis and our finding that PGCEs can predictably determine future evolutionary gains provide substantial evidence that the preponderance of parallel evolution over convergent evolution (Ord and Summers 2015; Conte et al. 2012) may be the result of specific, identifiable genetic dependencies entraining the evolutionary trajectory taken by similar genomes.

**Figure 5.**
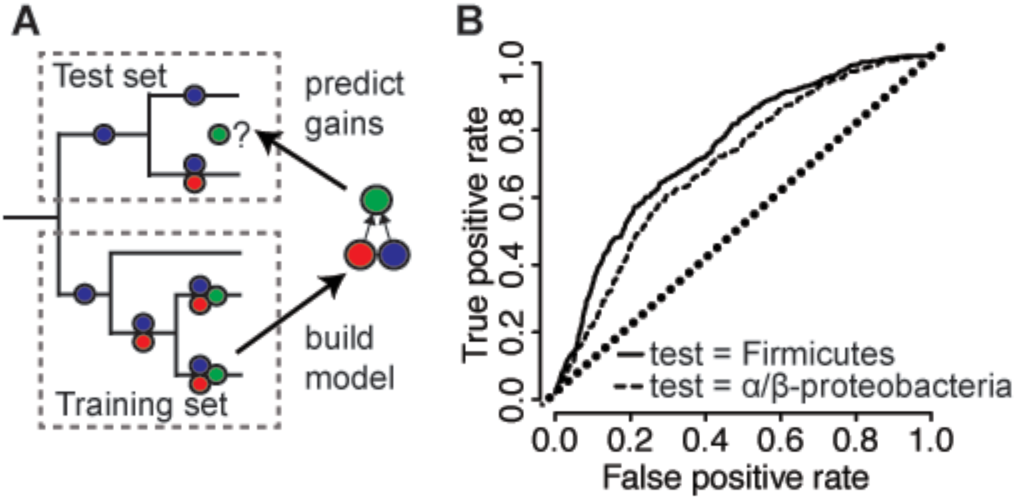
PGCE dependencies lead to taxonomically robust predictability of gene acquisition. (A): Workflow for predicting gene acquisition between clades of the tree. A training set is used to build a PGCE dependency model, which is then used to predict on which specific branches genes are likely to be gained (green circles), based on dependencies inferred from the training set (red and blue circles). (B): performance of PGCEs in predicting gene acquisitions in two test sets (indicated clades of the prokaryotic tree). Areas under each curve: Firmicutes, 0.73; Alpha/Beta-proteobacteria, 0.68. The diagonal dotted line represents the performance of a purely random prediction. See also Supplemental Figure S9.

## DISCUSSION

Combined, our findings provide substantial evidence to suggest that gene acquisitions in bacteria are governed by genome content through numerous gene-level dependencies. Our ability to detect these underlying dependencies is clearly imperfect, owing to various data and methodological limitations (Supplemental Text, Supplemental Figure S3). Therefore, in reality the complete dependency network is likely much denser than that described above and includes numerous dependencies and constraints that our approach may not be able to detect. Consequently, our estimates should be considered as a lower bound on the extent of gene-gene interactions, and accordingly, the predictability of HGT.

Notably, even considering such caveats, our observations dramatically expand our knowledge of the constraints on HGT. Previous studies of such constraints demonstrated that genes frequently acquired by HGT tend to occupy peripheral positions in biological networks, are often associated with specific cellular functions, and are phylogenetically clustered (Jain et al. 1999; Cohen et al. 2011). These observations suggested that properties of transferred genes are also important determinants of HGT regardless of recipient genome content (Jain et al. 1999; Cohen et al. 2011; Gophna and Ofran 2011) and that the acquisition of certain genes is clade-specific (Popa et al. 2011; Andam and Gogarten 2011). In contrast, our analysis demonstrates the importance of recipient genome content in influencing the propensity of a new gene to be acquired. In fact, to some extent, properties previously reported as determining the general “acquirability” of genes across all species may reflect an average constraint across genomes. By considering also variation in genomes acquiring genes, our analysis focused on specific biological effects, whose strengths may vary from genome to genome.

Importantly, our model that gene acquisition is affected by recipient genome content is consistent with the observed enrichment of HGT among close relatives, which presumably have similar genome content (Gogarten et al. 2002; Andam and Gogarten 2011; Popa et al. 2011; Popa and Dagan 2011). This taxonomic clustering of innovation by HGT is also in agreement with previous studies that demonstrated that phenotypic and genetic parallel evolution is more common than convergent evolution, potentially due to the effects of historical contingency (Gould and Lewontin 1979; Conte et al. 2012; Christin et al. 2015; Ord and Summers 2015). However, in contrast to other studies, we present direct evidence that the mechanism by which contingency controls evolution is epistasis. Furthermore, the universality of PGCEs shows that the constraints underlying the effect of contingency operate outside the context of parallel evolution.

Put differently, since each phylum-level clade is subject to an independent evolutionary trajectory, it is unlikely that the same dependency patterns would repeat solely due to parallel evolution. Moreover, our ability to predict where exactly along the tree gains of a specific gene are likely to occur (Figure 5B) suggests that PGCEs successfully capture how *variation* in the genomic content (even among closely related species) affects future gain events. Such PGCE specificity therefore indicates that observed dependencies are not a trivial byproduct of prevalent gene transfer events among taxonomically closely related genomes (e.g., due to homologous recombination constraints; Popa et al. 2011). Nonetheless, the relative contributions of each of these various processes governing the assembly of prokaryotic genomes (and the evolution of complex systems in general) clearly deserve future study.

It should also be noted that while our analysis revealed several intriguing patterns, the precise interpretation of some of these patterns remains unclear. For instance, the observed correspondence of topological ranks of genes to chronology suggests that evolutionary age is a potential contributor to such ranking, especially considering that our reconstructions likely lack many genes that have not been retained in any extant genomes. However, the biological plausibility and statistical robustness of PGCEs demonstrated above strongly argue that the observed evolutionary patterns are the result of constraint-inducing dependencies. Future work may therefore aim to quantify the trade-off between functional and chronological determinants in apparent evolutionary constraints.

Finally, we demonstrate the predictability of genomic evolution by horizontal transfer from current genomic content. As stated above, this finding also suggests that such dependencies are fairly universal across the prokaryotic tree. It should be noted that our approach was designed specifically to understand the PGCE network’s significance and universality, rather than predict gene acquisition. It is likely that an approach specifically engineered for gene acquisition prediction would substantially outperform our approach. The estimates of predictability of genomic evolution presented here are accordingly quite conservative.

The determinism and predictability of evolutionary patterns therefore appear to be an outcome not only of intramolecular epistasis in proteins or phylogenetic constraints, but also of genome-wide interactions between genes. This suggests that the evolution of medically, economically, and ecologically important traits in prokaryotes depends on ancestral genome content and is hence at least partly predictable, potentially informing research in the epidemiology of infectious diseases, bioengineering, and biotechnology.

## METHODS

All mathematical operations and statistical analyses were performed in R 2.15.3 (R Core Team 2016). Probabilistic ancestral reconstructions were obtained using the *gainLoss* program (Cohen and Pupko 2010). Phylogenetic simulations and plots were performed with the APE library (Paradis et al. 2004). Network analyses and algorithms were implemented using either the *igraph* (Csardi and Nepusz 2006) or *NetworkX* (Hagberg et al. 2013) libraries, and visualized using Cytoscape v3.1.1 (Shannon et al. 2003).

### Phylogenies

We used a pre-computed phylogenetic tree (Dehal et al. 2010) as a model of bacterial evolution. We mapped all extant organisms in this tree to organisms in the KEGG database by their NCBI genome identifiers, and pruned all tips that did not directly and uniquely map to KEGG. This yielded a phylogenetic tree connecting 634 prokaryotic species. For analyses involving subtrees of this phylogenetic tree, we used iTOL (Letunic and Bork 2011) to extract subtrees.

### Inferring phylogenetic histories for genes

We used the *gainLoss* v1.266 software (Cohen and Pupko 2010), a set of presence/absence patterns of orthologous genes from KEGG (Kanehisa et al. 2012), and the phylogenetic tree described above to infer 1) the probabilities of presence and absence of genes at internal nodes of the tree, 2) gain and loss rates of each gene, and 3) tree branch lengths within a single model. Specifically, in running *gainLoss*, we assumed a stationary evolutionary process, with gene gain and loss rates for each gene modeled as a mixture of three rates drawn from gamma distributions defined based on overall initial presence/absence patterns. A complete list of parameters used for *gainLoss* runs is given in the Supplemental Text and as Supplemental File S2. The *gainLoss* log file for the principal run on the full tree is also included as Supplemental File S3. Based on these models, we obtained a probabilistic ancestral reconstruction based on stochastic mapping for each of 5801 genes that were present in at least one species and absent in at least one species, and filtered out genes that were found to be gained less than twice throughout the tree, yielding 5031 genes which we further analyzed.

### Inferring gains and presence of genes on branches

To focus on gain events with strong support and where the gained gene is retained (rather than gain events where the gene is subsequently lost along the same branch), we used a simple model for computing the probability of different evolutionary gain/loss scenarios based on *gainLoss* ancestral reconstructions rather than directly using *gainLoss* gain inferences (Supplemental Text). Specifically, we assumed that unobserved gains and losses are not relevant, and that evolutionary scenarios are defined by the states at the ancestor and descendant nodes of each branch (regardless of branch length). With these assumptions, we used the probabilities of presence and absence of each of 5031 genes at each node and tip on the tree to compute the probability of each branch undergoing each scenario: 1) gain (absent in ancestor and present in descendant), 2) presence (present in both ancestor and descendant), and 3) loss (present in ancestor and absent in descendant; Supplemental Text). For a gene X on a branch with ancestor A and descendant B, we assume:

1. Pr(X*present* on branch) = Pr(X present in A ∩ X present in B) = Pr(X present in A) * Pr(X present in B)
2. Pr(X *gained* on branch) = Pr(X absent in A ∩ X present in B) = Pr(X absent in A) * Pr(X present in B)
3. Pr(X *lost* on branch) = Pr(X present in A ∩ X absent in B) = Pr(X present in A) * Pr(X absent in B)

Note again that these probability estimates are distinct from those obtained by using the *gainLoss* continuous-time Markov chain on the same ancestral reconstruction, which consider also hypothetical gains that are not retained and are thus not relevant to our analysis (Supplemental Text).

### Robustness analysis of reconstruction method

We used a maximum-parsimony reconstruction as inferred by *gainLoss* to benchmark the accuracy of the *gainLoss* reconstruction by stochastic mapping. In this analysis, only internal node reconstructions were considered, as tip reconstructions (for which the states are known) are not informative about algorithm performance. Since the maximum-parsimony reconstruction is binary (presence/absence) and the stochastic mapping reconstruction is probabilistic, for purposes of comparison we rounded the probabilities of the stochastic mapping reconstruction to obtain a presence/absence reconstruction (*i.e*., a probability >0.5 denotes presence and <=0.5 denotes absence). We computed the agreement between the two reconstructions as the percentage of internal node reconstructions that agree on the state of the gene.

### Comparison of analyzed gains to reconciliation-based HGT inference

We compared gains inferred by our method for several genes central to the PGCE network to gain events reported in a searchable database of horizontally acquired genes inferred by a sequence-based reconciliation method (Jeong et al. 2015). To this end, we classified all branches supporting a gain event for each of these genes with >50% probability by our method as *‘true ’* gains. We next searched the reconciliation database (all queries performed between January 15^th^ and February 20^th^, 2016) for each gene, identifying orthologous genes across 2,472 genomes that exhibit HGT according to reconciliation (excluding events that occurred on branches without descendants). We manually compared descendants of the remaining events from our method with the genomes experiencing gene acquisition in the reconciliation dataset to assess overlap between these two methods (see Supplemental Text).

### Quantifying PGCEs

We defined a “pair of genes with conjugated evolution” (PGCE) as a gene pair (*i, j*) for which the presence of one gene *i* encourages the gain of the other, *j*. Considering these genes as phylogenetic characters, we therefore aim to detect pairs for which “gain” state transitions for character *j* are enriched on branches where character *i* remains in the “present” state. This problem is related to previous methods for detecting coevolution or correlation between phylogenetic characters (Maddison 1990; Huelsenbeck et al. 2003; Cohen et al. 2012). Given *N* branches and *k* genes, there are 2 *N* X *k* matrices, *P* and *G*, describing the probabilities, respectively, of presence and gain of each gene along each branch (using our model for estimating gains described above). The test statistic for a dependency between each gene pair (*i, j*) is the expected number of branches where the gain of gene *j* occurs, while conditioning on the presence of gene *i* (cell *C_ij_* in a *k* x *k* matrix *C*). Counting transitions of one character (gene *j* gain) given some state of another character (gene *i* presence) yields a standard test statistic for testing correlated evolution of binary characters on phylogenies (Maddison 1990). To compute *C* across *N* branches, we sum the conditional probabilities of the gain of gene *j* in the presence of gene *i* across the tree, *i.e*. the products of the two *N*x *k* matrices, *P* (presence) and *G* (gain), for each gene pair:

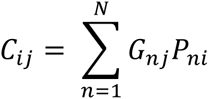

Entries in *C* which are significantly larger than a null expectation of gains represent PGCEs between the row and column genes of *C*.

### Null distribution for PGCEs

For two independently evolving genes *i* and *j*, the counted gains of *j* in the presence of *i, C_ij_*, will be distributed under the null hypothesis (independent evolution) as some function of the prevalence of *i* (the sum of *P_i_*, the vector of probabilities of presence of *i* across branches of the tree), the expected number of branches where *j* is gained (the sum of *G_j_*, the vector of probabilities of gains of *j* across nodes of the tree), and the topology and branch lengths of the tree (τ):

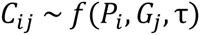

We followed previous studies (Cohen et al. 2012; Huelsenbeck et al. 2003; Maddison 1990) by approximating this null distribution via parametric bootstrapping. Specifically, we simulated the evolution of 10^5^ genes along the tree using the APE library function *rTraitDisc()* (Paradis et al. 2004). For the gain and loss rates used in these simulations, we used *gainLoss* gain and loss rates estimated for the 5801 empirical genes. We fit gamma distributions to these values by maximum likelihood using the function *fitdistr()* from the MASS library (Venables and Ripley 2002). For both gains and losses, we increased the shape parameter of the gamma distribution (by a factor of 3 for gains, 1.5 for losses), to ensure that simulated genes showed sufficiently large numbers of gains. This was necessary because parametric bootstrapping with the rates inferred by *gainLoss* resulted in left skewed distributions of gene gains (compare Supplemental Figures S2A, S2C, and S2E), which were likely to confound null models. For our null models to be applicable, the distribution of simulated gene gains should be roughly similar to the distribution of gains among empirical genes (see Supplemental Figure S2, Supplemental Text).

These simulated genes should evolve independently and thus represent a null model for PGCEs. As above, we constructed matrices representing the probabilities of presence and gain of these 10^5^ genes across all of the branches of the phylogeny (*P_null_* and *G_null_*). We then multiplied these matrices of simulated genes to compute a 10^5^ x 10^5^ matrix *C_null_* of expected branch counts under a model of independence. We excluded gene pairs with *C_ij_* ≤ 1 from further analysis, as it may be difficult to distinguish between no association and a lack of statistical power for such pairs (Supplemental Figure S3A), reducing overall power in computing false discovery rates (Bourgon et al. 2010). As a null distribution for each pair of genes *i* and *j* with *C_ij_* > 1, we used the 1000 simulated genes with prevalence closest to gene *i* (rows of *C_null_*), and the 1000 simulated genes with a number of gains closest to gene *j* (columns of *C_null_*). We used the 10^6^ simulated observations in the resulting submatrix of *C_null_* as a null distribution for *C_ij_*. Notably, *C_ij_* includes non-integer count expectations, whereas *C_null_* represents integer counts (because the true reconstruction is known). Consequently, we floored values in *C_ij_*, such that all counts were truncated at the decimal point. The comparison of *C_ij_* to this null distribution yields an empirical p-value; we rejected the null hypothesis of independence between genes *i* and *j* for the *C_ij_* observation at a 1% false discovery rate (Benjamini and Hochberg 1995) (P < 7 x 10^−6^).

### Constructing a PGCE network

For each entry in *C_ij_* for which we observed a significant association, we recorded an edge from gene *i* to gene *j* in a network of PGCEs. To focus purely on direct interactions, we subjected this network to a transitive reduction (Hsu 1975). This reduction requires a directed acyclic graph (DAG). To identify the largest possible DAG in our PGCE network, we identified and removed the minimal set of edges inducing cycles (Supplemental Text). We performed a transitive reduction of the resulting DAG using Hsu’s algorithm (Hsu 1975) (Supplemental Text).

### Mapping biological information to the network

We used network rewiring (as implemented in the *rewire()* function of the *igraph* library (Csardi and Nepusz 2006)) to generate null distributions of the PGCE network by randomly exchanging edges between pairs of connected nodes, while excluding self-edges. In each permutation, we performed 5N rewiring operations, where there are N edges in the network, to ensure sufficient randomization. To estimate the relationship between the PGCE network and biological information we calculated the number of edges shared between the PGCE network and a metabolic network of all bacterial metabolism obtained from KEGG (Kanehisa et al. 2012; Levy and Borenstein 2013), and the number of edges shared between members of the same functional pathway as defined by KEGG, in both the original and randomized networks.

To determine whether genes with certain functional annotations were more likely to associate with one another in the PGCE network, we examined the KEGG Pathway annotations of each pair of genes in the network. We counted the number of edges leading from each pathway to each other pathway, and obtained an empirical p-value for this count by comparing it to a null distribution of the expected counts obtained by random rewiring as above.

### Topological sorting of PGCE networks

To identify global patterns in our PGCE network, we performed topological sorting (Kahn 1962) with grouping. Topological sorting finds an absolute ordering of nodes in a directed acyclic graph (DAG), such that no node later in the ordering has an edge directed towards a node earlier in the ordering. Grouping the sort allows nodes to have the same rank in the ordering if precedence cannot be established between them, giving a unique solution. For a description of the algorithm used, see Supplementary Text.

### Prediction of HGT events on branches

We used the PGCE network to predict the occurrence of specific HGT events (gene acquisitions) on the tree in the following fashion. We used two test/training set partitions, with the clades of Firmicutes and the Alpha/Betaproteobacteria as independent test sets, and the training sets as the rest of the tree without these clades. To “train” PGCE networks, we performed ancestral reconstruction of gene presence, PGCE inference, and network processing just as for the entire tree. We only attempted to predict genes with at least one PGCE dependency (“predictable” genes). We then considered each branch in the test set independently, attempting to predict whether each predictable gene was gained on that branch based on the reconstructed genome at the ancestor node. For each predictable gene-branch combination, our prediction score was the proportion of the predictable gene’s PGCE dependencies that are present in the ancestor. This is the dot product of the gene presence/absence pattern of the ancestor node (*A_i_* across *i* potentially present genes) and a binary vector denoting which genes in the PGCE network the predictable gene depends on (*P_i_* across *i* genes in potential PGCEs), scaled by *P_i_*:

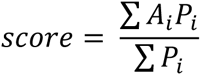

Note that this value ranges between 0 and 1 for each predicted gene. As true gains, we used our reconstructed gene acquisition events for each branch in the test set. We arbitrarily called any predictable gene-branch pair with a Pr(gain) > 0.5 as a gain, and any predictable gene-branch pair with Pr(gain) <= 0.5 as no gain. We filtered out any gene-branch pair where the gene was known to be present with Pr > 0.4, as in these cases the gene is probably already present. We analyzed the accuracy of our prediction scores using receiver operating characteristic (ROC) analysis and by comparing scores of the gain branches to those of the no-gain branches.

## Data Access

Parameter and log files for principal analyses are provided as Supplemental Files S2 and S3. Data and code are provided as Supplemental File S4.

## Acknowledgements

We are obliged to members of the Borenstein and Queitsch laboratories, and to Evgeny Sokurenko, Joe Felsenstein, and Willie Swanson for helpful discussions. We thank Ofir Cohen for help with the *gainLoss* program. We thank Hyeon Soo Jeong for help with the HGTree database. MOP was supported in part by National Human Genome Research Institute Interdisciplinary Training in Genome Sciences Grant 2T32HG35-16. CQ is supported by National Institute of Health New Innovator Award DP2OD008371. EB is supported by National Institute of Health New Innovator Award DP2AT00780201. We thank UW Genome Sciences Information Technology Services for high-performance computing resources.

## REFERENCES

Andam CP, Gogarten JP. 2011. Biased gene transfer in microbial evolution. Nat Rev Microbiol 9: 543–55.

Andersson I, Backlund A. 2008. Structure and function of Rubisco. Plant Physiol Biochem 46: 275–91.

Baltrus DA. 2013. Exploring the costs of horizontal gene transfer. Trends EcolEvol 28: 489–95.

Benjamini Y, Hochberg Y. 1995. Controlling the false discovery rate: a practical and powerful approach to multiple testing. JR Stat Soc B 57: 289–300.

Bourgon R, Gentleman R, Huber W. 2010. Independent filtering increases detection power for high-throughput experiments. Proc Natl Acad Sci U S A 107: 9546–51.

Chen HD, Jewett MW, Groisman E a. 2011. Ancestral genes can control the ability of horizontally acquired loci to confer new traits. PLoS Genet 7: e1002184.

Christin P-A, Arakaki M, Osborne CP, Edwards EJ. 2015. Genetic enablers underlying the clustered evolutionary origins of C4 photosynthesis in angiosperms. Mol Biol Evol 32: 846–58.

Cohen O, Ashkenazy H, Burstein D, Pupko T. 2012. Uncovering the co-evolutionary network among prokaryotic genes. Bioinformatics 28: i389–i394.

Cohen O, Gophna U, Pupko T. 2011. The complexity hypothesis revisited: connectivity rather than function constitutes a barrier to horizontal gene transfer. Mol Biol Evol 28: 1481–9.

Cohen O, Pupko T. 2010. Inference and characterization of horizontally transferred gene families using stochastic mapping. Mol Biol Evol 27: 703–13.

Conte GL, Arnegard ME, Peichel CL, Schluter D. 2012. The probability of genetic parallelism and convergence in natural populations. Proc R Soc B Biol Sci 279: 5039–47.

Csardi G, Nepusz T. 2006. The igraph Software Package for Complex Network Research. InterJournal Complex Sy.

Dehal PS, Joachimiak MP, Price MN, Bates JT, Baumohl JK, Chivian D, Friedland GD, Huang KH, Keller K, Novichkov PS, et al. 2010. MicrobesOnline: an integrated portal for comparative and functional genomics. Nucleic Acids Res 38: D396–400.

Delwiche CF, Palmer JD. 1996. Rampant horizontal transfer and duplication of rubisco genes in eubacteria and plastids. Mol Biol Evol 13: 873–882.

Eppley RW, Peterson BJ. 1979. Particulate organic matter flux and planktonic new production in the deep ocean. Nature 282: 677–680.

Gogarten JP, Doolittle WF, Lawrence JG. 2002. Prokaryotic evolution in light of gene transfer. Mol Biol Evol 19: 2226–38.

Gong LI, Suchard MA, Bloom JD. 2013. Stability-mediated epistasis constrains the evolution of an influenza protein. Elife 2: e00631.

Gophna U, Ofran Y. 2011. Lateral acquisition of genes is affected by the friendliness of their products. Proc Natl Acad Sci U S A 108: 343–8.

Gould SJ, Lewontin RC. 1979. The Spandrels of San Marco and the Panglossian Paradigm: A Critique of the Adaptationist Programme. Proc R Soc B Biol Sci 205: 581–598.

Hagberg A, Schult D, Swart P. 2013. NetworkX. High productivity software for complex networks. https://networkx.lanl.gov/.

Harms MJ, Thornton JW. 2014. Historical contingency and its biophysical basis in glucocorticoid receptor evolution. Nature 512: 203–7.

Hsu HT. 1975. An Algorithm for Finding a Minimal Equivalent Graph of a Digraph. J ACM 22: 11–16.

Huelsenbeck JP, Nielsen R, Bollback JP. 2003. Stochastic Mapping of Morphological Characters. Syst Biol 52: 131–158.

Iwasaki W, Takagi T. 2009. Rapid pathway evolution facilitated by horizontal gene transfers across prokaryotic lineages. ed. I. Matic. PLoS Genet 5: e1000402.

Jain R, Rivera MC, Lake J a. 1999. Horizontal gene transfer among genomes: the complexity hypothesis. Proc Natl Acad Sci U S A 96: 3801–6.

Jain R, Rivera MC, Moore JE, Lake JA. 2003. Horizontal gene transfer accelerates genome innovation and evolution. Mol Biol Evol 20: 1598–602.

Jeong H, Sung S, Kwon T, Seo M, Caetano-Anollés K, Choi SH, Cho S, Nasir A, Kim H. 2015. HGTree: database of horizontally transferred genes determined by tree reconciliation. Nucleic Acids Res gkv1245–.

Johnson CM, Grossman AD. 2014. Identification of host genes that affect acquisition of an integrative and conjugative element in Bacillus subtilis. Mol Microbiol.

Kahn AB. 1962. Topological Sorting of Large Networks. Commun ACM 5: 558–562.

Kanehisa M, Goto S, Sato Y, Furumichi M, Tanabe M. 2012. KEGG for integration and interpretation of large-scale molecular data sets. Nucleic Acids Res 40: D109–14.

Lerat E, Daubin V, Ochman H, Moran NA. 2005. Evolutionary origins of genomic repertoires in bacteria. ed. D. Hillis. PLoS Biol 3: e130.

Letunic I, Bork P. 2011. Interactive Tree Of Life v2: online annotation and display of phylogenetic trees made easy. Nucleic Acids Res 39: W475–8.

Levy R, Borenstein E. 2013. Metabolic modeling of species interaction in the human microbiome elucidates community-level assembly rules. Proc Natl Acad Sci USA 110: 12804–9.

Maddison WP. 1990. A Method for Testing the Correlated Evolution of Two Binary Characters: Are Gains or Losses Concentrated on Certain Branches of a Phylogenetic Tree? Evolution (N Y) 44: 539–557.

Mira A, Ochman H, Moran NA. 2001. Deletional bias and the evolution of bacterial genomes. Trends Genet 17: 589–596.

Nowell RW, Green S, Laue BE, Sharp PM. 2014. The extent of genome flux and its role in the differentiation of bacterial lineages. Genome Biol Evol 6: 1514–29.

Ord TJ, Summers TC. 2015. Repeated evolution and the impact of evolutionary history on adaptation. BMC Evol Biol 15: 137.

Orr HA. 2005. The Probability of Parallel Evolution. Evolution (N Y) 59: 216–220.

Pal C, Papp B, Lercher MJ. 2005. Horizontal gene transfer depends on gene content of the host. Bioinformatics 21: ii222–ii223.

Paradis E, Claude J, Strimmer K. 2004. APE: Analyses of Phylogenetics and Evolution in R language. Bioinformatics 20: 289–290.

Van Passel MWJ, Marri PR, Ochman H. 2008. The emergence and fate of horizontally acquired genes in Escherichia coli. PLoS Comput Biol 4: e1000059.

Popa O, Dagan T. 2011. Trends and barriers to lateral gene transfer in prokaryotes. Curr Opin Microbiol 14: 615–23.

Popa O, Hazkani-Covo E, Landan G, Martin W, Dagan T. 2011. Directed networks reveal genomic barriers and DNA repair bypasses to lateral gene transfer among prokaryotes. Genome Res 21: 599–609.

Press MO, Li H, Creanza N, Kramer G, Queitsch C, Sourjik V, Borenstein E. 2013. Genome-scale co-evolutionary inference identifies functions and clients of bacterial Hsp90. PLoS Genet 9: e1003631.

Puigbò P, Lobkovsky AE, Kristensen DM, Wolf YI, Koonin E V. 2014. Genomes in turmoil: Quantification of genome dynamics in prokaryote supergenomes. BMC Biol 12: 66.

R Core Team. 2016. R: A Language and Environment for Statistical Computing. R Foundation for Statistical Computing, Vienna, Austria. http://www.R-project.org/.

Shannon P, Markiel A, Ozier O, Baliga NS, Wang JT, Ramage D, Amin N, Schwikowski B, Ideker T. 2003. Cytoscape: a software environment for integrated models of biomolecular interaction networks. Genome Res 13: 2498–504.

Smillie CS, Smith MB, Friedman J, Cordero OX, David L a, Alm EJ. 2011. Ecology drives a global network of gene exchange connecting the human microbiome. Nature 480: 241–4.

Sohm JA, Webb EA, Capone DG. 2011. Emerging patterns of marine nitrogen fixation. Nat Rev Microbiol 9: 499–508.

Sorek R, Zhu Y, Creevey CJ, Francino MP, Bork P, Rubin EM. 2007. Genome-wide experimental determination of barriers to horizontal gene transfer. Science 318: 1449–52.

Soucy SM, Huang J, Gogarten JP. 2015. Horizontal gene transfer: building the web of life. Nat Rev Genet 16: 472–482.

Venables WN, Ripley BD. 2002. Modern Applied Statistics with S. Fourth Edi. Springer, Springer.

De Visser JAGM, Krug J. 2014. Empirical fitness landscapes and the predictability of evolution. Nat Rev Genet 15: 480–490.

Weinreich DM, Delaney NF, Depristo MA, Hartl DL. 2006. Darwinian evolution can follow only very few mutational paths to fitter proteins. Science 312: 111–4.

